# Adult corals enhance juvenile recruitment through cumulative ecological niche construction

**DOI:** 10.64898/2026.09.21.753173

**Authors:** Viviana Brambilla, Mollie Asbury, Andrew H Baird, Matthew-James Bennett, Nader Boutros, James Cant, Cher F. Y. Chow, Garrett J. Fundakowski, Wilhelm Marais, Oscar Pizarro, Damaris Torres-Pulliza, Nina Schiettekatte, Rachael Woods, Stefan Williams, Devynn Wulstein, Kyle Zawada, Joshua Madin, Maria Dornelas

## Abstract

Ecosystem engineers are ecological niche constructors: by modifying local environments, they alter the conditions under which organisms establish, persist and interact. Although niche construction is often framed as a population-level evolutionary feedback, it could also operate at the assemblage level when communities collectively shape their environments and filter future trajectories, such as recruitment patterns. Reef-building corals offer an ideal setting to test this idea because adult colonies form three-dimensional carbonate structures that can persist after colony death, creating an inherited habitat for subsequent coral generations. Here, we tested whether coral recruitment to the juvenile stage was associated with local adult coral abundance, reef structure, and larval settlement supply. Combining spatially explicit coral censuses across >1300 quadrats, photogrammetric reef digital elevation models and data from settlement tiles across 15 reef sites at two time points at Jiigurru, Great Barrier Reef, we found that juvenile abundance increased additively on the log scale with adult abundance and with fractal dimension, a metric of fine-scale structural complexity. Surface rugosity had weaker effects, and interactions between adult abundance and habitat structure were limited, indicating largely additive, rather than synergistic or antagonistic, effects. Larval supply was negatively correlated with juvenile abundance. Our results show that both adult assemblages and the concurrent structural complexity, consistent with assemblage-level niche construction and ecological inheritance theory, shape coral recruitment.

## Introduction

Ecosystem engineers are ecological niche constructors: organisms whose presence and activities modify the local environment, affecting the rates at which organisms colonise, persist and interact ^1,2^. Although niche construction theory has often emphasised individual population evolutionary feedbacks, i.e. the selective advantage upon the population modifying the environment ^3,4^, the same logic can apply at the scale of ecological assemblages, where collectively constructed environments influence community composition, diversity and ecosystem processes ^5^. As in population genetics, community dynamics are organised around selection, drift, speciation, and dispersal, with individuals in assemblages playing a role analogous to alleles in populations: their relative species abundance changes through deterministic (selection) and stochastic (drift) processes, while new species enter through dispersal or speciation ^6^. From this perspective, niche construction becomes a community-level eco-evolutionary process whenever organisms alter the selective environment experienced by the rest of the community. Here, we test whether assemblage-level ecological niche construction affects the recruitment of scleractinian corals, a group of ecosystem engineers that act as foundation species for some of the most diverse ecosystems on the planet.

In diverse ecosystems, niche construction effects are rarely attributable to individual species of engineers; they can emerge instead from functional groups of species acting collectively ^5^. In tundra plant communities, forbs and grasses can be rare. Yet, as a group, they can strongly predict local plant species diversity, suggesting that less abundant species can nevertheless disproportionately structure the realised niches of co-occurring species ^5,7^. This raises a central question regarding whether assemblage-level niche construction effects are best detected from constructor abundance or density, from the magnitude and stability of their environmental modification, or from the interaction between the two. Density-dependent and environmental modification effects can be additive, antagonistic or synergistic, producing assemblage outcomes that cannot be inferred from abundance or engineering function alone (Fig. 1).

**Figure 1.**
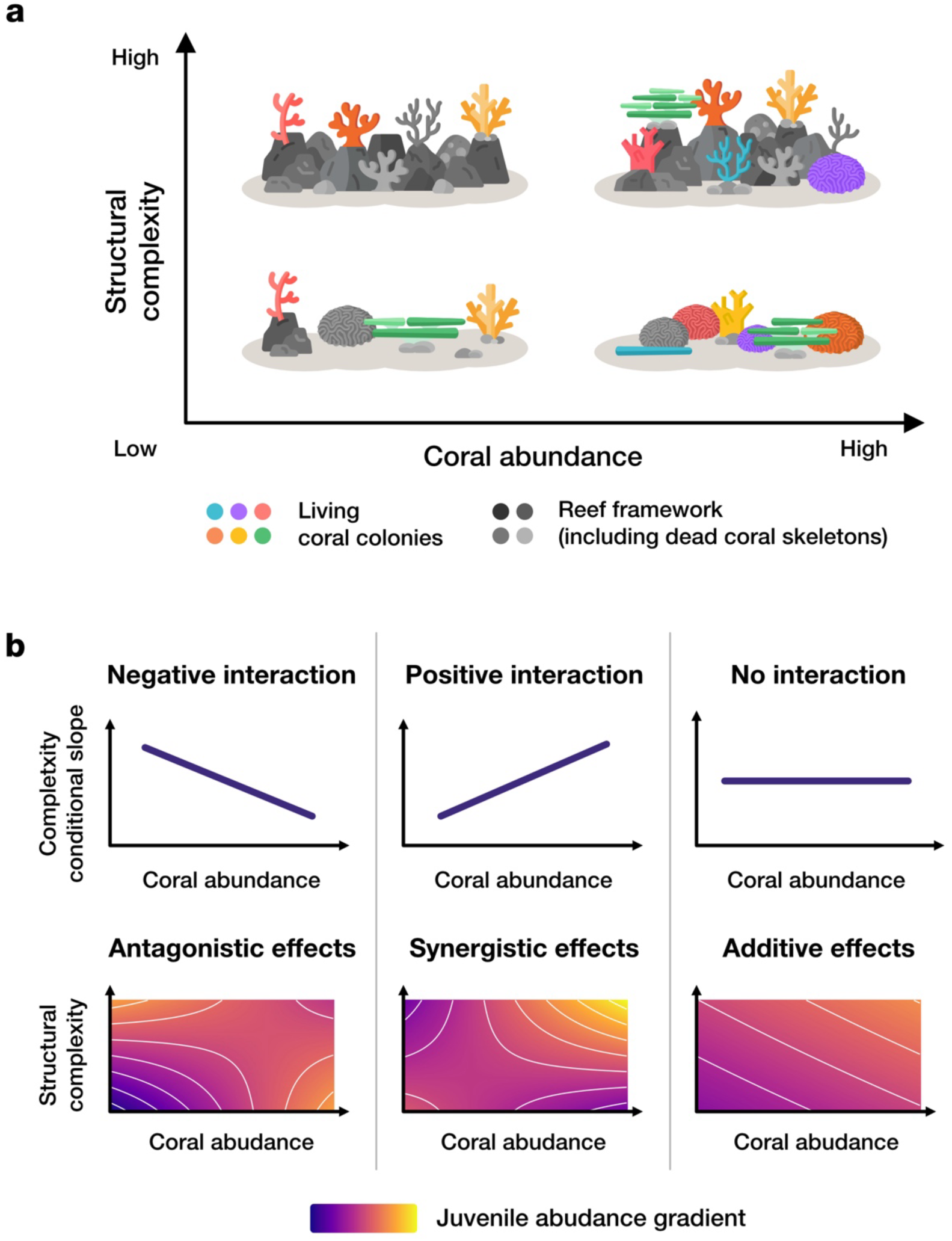
Reef structural complexity and adult abundance can have antagonistic, synergistic, or additive cumulative effects on juvenile recruitment. (a) Habitat structural complexity emerges from the reef framework as a whole and can therefore be decoupled from adult coral abundance. The space defined by structural complexity and adult abundance represents a continuum of reef configurations, each of which could have different consequences for juvenile recruitment. In the scheme, vignettes show higher complexity on the top row for fixed numbers of coral colonies (3 living colonies on the left, and 6 on the right) (b) When structural complexity and adult abundance are viewed together, their combined effects on juvenile abundance can be additive, antagonistic or synergistic. A negative interaction indicates that the effect of structural complexity weakens as adult abundance increases, producing an antagonistic effect and a shift in the direction of the juvenile abundance gradient across the complexity–abundance continuum. A positive interaction indicates that the effect of structural complexity strengthens with increasing adult abundance, producing a synergistic effect and a steeper juvenile abundance gradient along the complexity axis at high coral abundance. In the absence of an interaction, the effects are additive: structural complexity and adult abundance contribute independently to juvenile abundance, and the gradient direction remains constant across the continuum. Icons were inspired by CC0 icons deposited on Flaticon.com.

Density-dependent processes shape recruitment, since the probability that a juvenile recruits successfully often depends on the abundance, identity and proximity of adults already present ^8,9^. In sessile assemblages, abundant adults can, for example, reduce the survival of nearby recruits through competition for light, nutrients or space^9,10^, or through the accumulation of host-specific ‘enemies’ such as pathogens, herbivores or seed predators^11,12^. The Janzen–Connell effect is a prime example: recruit survival is lower near related, or more broadly conspecific, adults because these adults concentrate ‘enemies’ or deplete resources, disproportionately affecting similar juveniles^11,12^. Yet this effect can change when the relevant neighbourhood is defined not by close relatives or conspecifics, but by the adult assemblage as a whole. If recruits occur more often near unrelated adults, this positive association might arise because adults and juveniles respond to the same abiotic conditions and the environment that benefits adult persistence also filters for juveniles with similar requirements^13^. Alternatively, adult assemblages might actively facilitate recruitment by reducing stress^14,15^, stabilising soils^16^, trapping propagules^17^, or buffering temperature extremes^14,18^. Thus, adult abundance can generate contrasting recruitment signals depending on whether adults are treated as competitors, ‘enemy’ reservoirs, environmental indicators, facilitators or a mix of all these options.

Ecosystem engineers can provide highly complex three-dimensional habitats ^19^, which can positively influence biodiversity and ecosystem functioning. In terrestrial systems, forest structural complexity is associated with the diversity of multiple taxa, including birds ^20^ and arthropods ^21^, and with ecosystem productivity ^22^. These relationships reflect direct effects of structure, such as increasing the amount of surface or space available for colonisation, as well as indirect effects, because physical structures can modify microhabitats, abiotic conditions and trophic interactions ^19,23^. In shallow tropical waters, reef-building hard corals are a clear case of ecosystem engineers. By producing carbonate skeletons, corals generate habitat structure that changes the geometry of available space ^24^ and alters local environmental conditions^25^. These effects also persist beyond the lifespan of individual colonies as coral skeletons remain within the reef framework after colony death. This creates ecological inheritance through which past engineering shapes current habitats ^4^. As such, coral abundance can be decoupled from reef complexity (Fig. 1). Therefore, reef structural complexity partly reflects the cumulative and persistent effects of coral niche construction, i.e. how coral assemblages have modified, and continue to modify, the environment experienced by other organisms and themselves, including future coral recruits.

Corals have a planktonic larval stage, after which larvae settle on suitable substrata, metamorphose and begin their sessile life. As coral larvae are poor swimmers, they rely on small structural features that influence larval settlement by modifying near-reef flow ^26^. Successful recruitment depends on sufficient larval supply and settlement, but also on post-settlement mortality, which remains high during the first years as settled corals grow into juveniles ^10,27–30^. During this vulnerable period, the local community can strongly shape settlers’ survival through complex interactions that are at times positive and negative ^31,32^. For example, free substratum is necessary for growth, so high living coral cover can limit growth. Algae can further reduce settlement and juvenile survival by occupying substrata and limiting light ^33^. In contrast, herbivores can indirectly facilitate corals by controlling algae but can also damage small recruits while grazing ^27^. Habitat structural complexity can mediate many of these processes. Fine-scale refuges such as crevices can protect settler recruits and are often associated with higher juvenile abundance ^27,34^. However, complexity can also reduce light, trap sediment or alter exposure to grazers and predators. Therefore, settlement patterns alone might not predict recruitment outcomes. To understand which reef configurations support coral replenishment, it is essential to examine the relationships among adult corals, habitat structure and juvenile abundance after post-settlement mortality has occurred.

Reef complexity can influence benthic communities both directly, by providing surface area and spatial heterogeneity, and indirectly by modifying flow ^35^, sedimentation ^36^, light exposure ^37^, or predator–prey interactions ^38,39^. Yet one assemblage-level feedback remains unexplored: adult corals contribute to structural complexity, but that same complexity might in turn modify how adult abundance affects successful recruitment, defined here as the establishment of settlers into the juvenile stage. If adults reduce recruitment through competition, enemy accumulation, or space pre-emption, structural complexity could buffer negative density dependence by increasing refuge availability, expanding settlement surfaces, or modifying local environmental conditions. Conversely, if adults facilitate recruits, for example, by horizontal transmission of the symbionts^40^ or, more generally, their microbiome^41^, complexity might amplify the positive effects of adult abundance. Disentangling these scenarios requires asking whether recruitment is best predicted by adult abundance, habitat complexity, or their interaction. This distinction matters because additive, antagonistic, or synergistic effects between adult assemblages and reef structure would imply different niche-construction mechanisms regulating coral community dynamics. A positive interaction would indicate that the effect of one predictor becomes stronger as the other increases, suggesting synergistic effects. A negative interaction would indicate that the effect of one predictor weakens as the other increases, consistent with antagonistic or buffering effects. If no interaction is detected, but both predictors have significant individual effects, this would suggest that adult abundance and structural complexity affect recruitment additively and independently.

Here, we investigate how density-dependent processes and organism-mediated environmental modification jointly shape successful recruitment in a natural shallow-reef coral assemblage. Specifically, we tested the effects of adult coral abundance and reef structural complexity, quantified using fractal dimension and surface rugosity, on juvenile coral abundance in reef patches at Jiigurru, Great Barrier Reef. Because reef complexity can be produced by adult coral assemblages and might therefore covary with adult abundance, we tested whether adult abundance and structural complexity acted independently or interactively to influence juvenile abundance. We predicted that sites with greater larval supply would have higher juvenile abundance. We also expected fractal dimension to be positively associated with juvenile abundance, potentially because complex reef structures can generate micro-eddies that enhance larval retention and settlement. Surface rugosity was also expected to promote recruitment by increasing settlement surface area and environmental heterogeneity per unit planar area. Our results show that recruitment increases with adult abundance but also additively depends on reef structural configurations, indicating that density-dependent processes and different dimensions of habitat complexity jointly shape early coral community assembly.

## Methods

Data was collected as part of a yearly spatially explicit monitoring project of coral communities, and focused on 15 reef sites surrounding Jiigurru (aka Lizard Island) in 2019 and 2023 ^42^. Annually since 2018, digital elevation models (DEMs, 5mm/pixel) and orthomosaics representing 120-square-meter reef sites were obtained by processing photos captured with the spiral method using Agisoft Metashape ^42–44^. Each year, locations of coral colonies were annotated in-water on orthomosaics printed on underwater paper, and the status (alive, new, dead) of every colony was confirmed across consecutive years. Cryptic colonies, that is, colonies that were visible in the field but not on the orthomosaics because of their size or their position, were also tracked. The geographic coordinates of each coral colony were obtained by digitising field annotations on the georeferenced reef orthomosaics in QGIS ^45^. Juveniles were defined as new coral colonies, and adults as previously recorded coral colonies that were still alive at the time of annotations. Sites and years for the analysis were selected to ensure the same time window, i.e., 12 months, across annotations.

To quantify the effect of spatial scale, sites were subdivided into different sized patches (Figs. SM1 and SM2). An 8 x 8 m square was centred in each of the reef DEMs and divided into grids with patch dimensions of 0.5 x 0.5 m, 1 x 1 m, and 2 × 2 m. This process resulted in 256, 64, and 16 reef patches of 0.25 m^2^, 1 m^2^, and 4 m^2^ respectively, per site per year. The primary analysis was performed using the 1 x 1 m cell size, and the other two scales (0.5 x 0.5 m and 2 x 2 m) were used to test whether observed relationships were robust across scales. DEMs for each site were cropped for each patch to calculate surface rugosity and fractal dimension (Fig. SM1). These metrics were chosen because they independently capture different aspects of complexity ^42,46^ and are common metrics in reef ecology ^47,48^. Surface rugosity and fractal dimension were obtained with the ‘hdr’ function of the package ‘habtools’ ^46^, specifying the ‘area’ method for surface rugosity and the ‘var’ method for fractal dimension ^46^, and using a 5 mm resolution. Juvenile and adult counts for each patch were extracted from the annotation layer using the ‘sf’ ^49^ and the ‘terra’ ^50^ packages.

To assess settler supply, the abundance of new coral settlers from the previous spawning events was estimated using settlement tiles^51^. At every site, six settlement tiles (11 × 11 cm, unglazed clay) were attached to the reef at least 3 m apart from each other, 5 days before the predicted spawning date ^52^. Eight weeks after spawning, tiles were retrieved (January 2019 and 2023), soaked in a mix of water and bleach, and then left to dry. Each tile was carefully inspected under a stereo-dissecting microscope, and all coral settlers were counted. The average tile count of coral settlers per site was used as a measure of coral settlement during the spawning events before our juvenile surveys took place.

Bayesian generalized mixed effect models were fitted for each scale to examine the effects of patch-level adult abundance, patch-level reef complexity, and site-level settlement on juvenile abundance in each grid cell. In each patch, juvenile abundance was modelled as a function of adult abundance, average settler abundance, surface rugosity, and fractal dimension, allowing each habitat and settler predictor to interact with adult abundance. To account for spatial variation across sites and temporal variation across years, the models included site and year as random effects. When appropriate, predictors were transformed to improve their distribution and range (Surface rugosity was log10-transformed, adult and settlement abundances were log-transformed), and all variables were centred to zero to ease interpretation. Because the error distribution exceeded the mean, appropriate error distributions were chosen for each grid size: a negative binomial distribution with a log link function. All the priors for model fitting were left as defaults (flat priors), and for each model four chains for 10000 iterations were run, with a warm-up period of 2000 iterations and a thinning rate of 5 iterations. We ensured that Rhat values were ∼ 1, and goodness of fit was assessed by visual inspection of the chains and the posterior predictive plots. Models were fitted with the probabilistic language RStan using the ‘brms’ package ^53^ and graphic representations of their output produced with ‘tidybayes’ ^54^, and ‘ggplot2’ ^55^. All analyses were performed in R^56^.

## Results

We recorded 22,763 corals across the two survey years, comprising 14,036 adults and 8,727 juveniles. The primary dataset (1 × 1 m scale) comprised 1,344 quadrats distributed across 15 sites (Fig. 2). Adult and juvenile abundances were right-skewed, with many cells containing relatively few individuals and a smaller number of high-count cells. At the 1 m scale, adult abundance ranged from 0 to 53 individuals per cell, with a median of 8 (interquartile range = 4–15), whereas juvenile abundance ranged from 0 to 44 individuals per cell, with a median of 4 (interquartile range = 1–10). Mean settlement varied from 0.5 to 25.8, with a median of 6.3. Habitat-structure variables also spanned broad ranges: surface rugosity ranged from 1.06 to 5.15, and fractal dimension from 1.80 to 2.47 (Fig. 2). Fractal dimension values below 2 are expected as an artefact of the implemented variation method. The datasets for scale sensitivity analysis retained the same site and year structure but differed in spatial grain, comprising 5,376 cells at 0.5 x 0.5 m scale (Fig. SM4) and 336 cells at 2 x 2 m scale (Fig. SM5).

**Figure 2.**
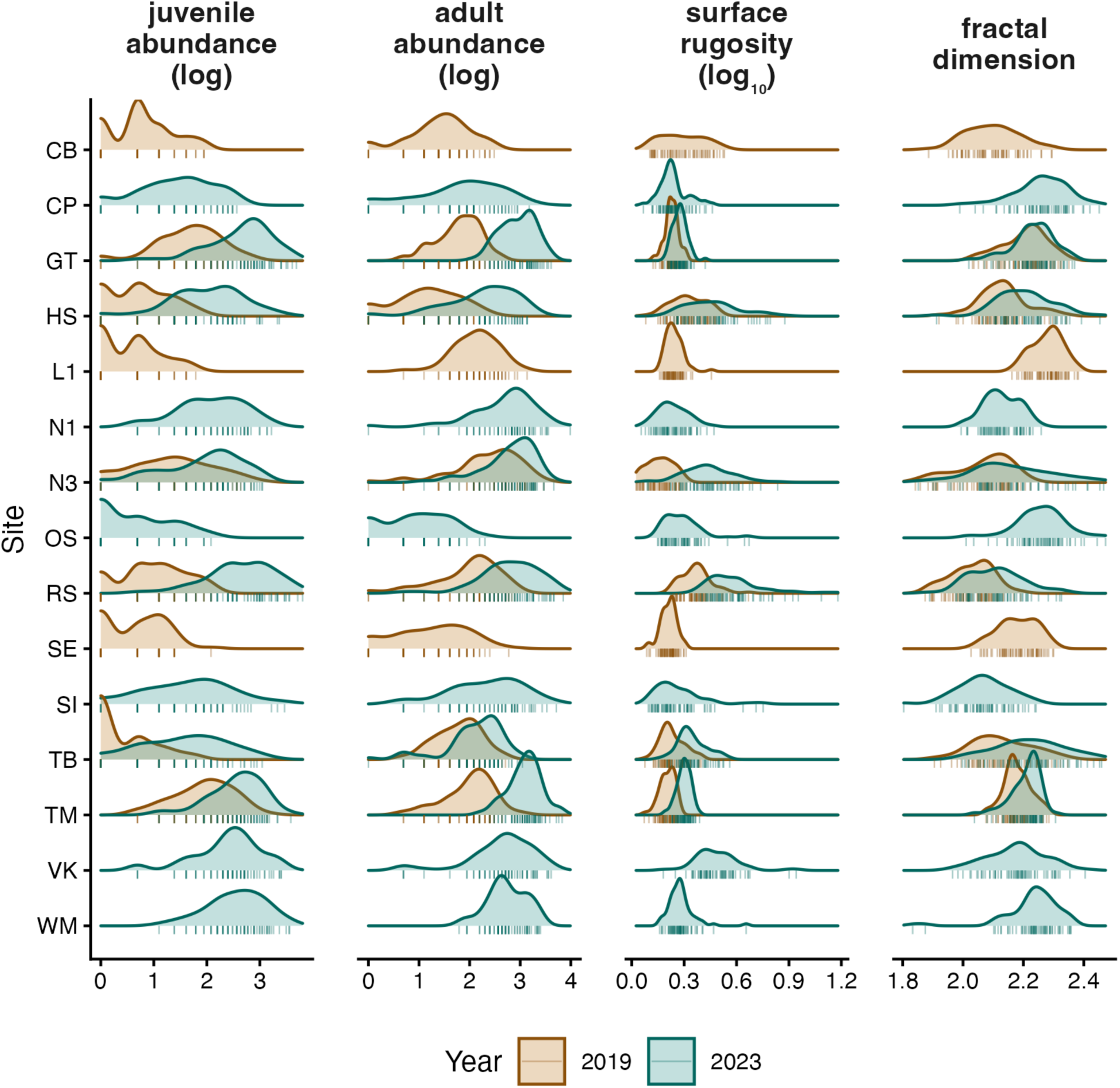
Density distributions of the variables for the primary dataset (1 m × 1 m scale) show overall variation across sites and years sampled (n = 1344). (a) Juvenile abundances were right-skewed, particularly in 2019, with many patches containing relatively few individuals and a smaller number of high abundance patches. (b) Adult abundances were less right-skewed, and in sites with repeated samples, they were higher in 2023. (c) Surface rugosity varied by an order of magnitude and reached very high values, particularly in deeper sites. (d) Fractal dimension was normally distributed and remained quite consistent across years. For comparisons with other scales, refer to Figure SM4 and SM5.

None of the three interactions between adult abundance and the habitat or settler predictors were credibly different from zero (adult × settlers: posterior mean = 0.02, 95% CI = [−0.06, 0.10]; adult × rugosity: posterior mean = −0.22, 95% CI = [−0.64, 0.21]; adult × fractal dimension: posterior mean = −0.19, 95% CI = [−0.77, 0.40], Figs. 3 and SM6). Examining the conditional slopes across the observed range of adult abundance corroborates this interpretation (Figs. 3b and SM7). The slope of average settlers on juvenile abundance remained essentially constant and negative across all levels of adult abundance, with a 95% credible band that did not include zero at any point in the range. The slope of fractal dimension was positive across the entire observed range of adult abundance, with the median slope decreasing modestly from ∼1.1 at low adult densities to ∼0.4 at high adult densities; however, the credible band widened substantially at the extremes and overlapped zero only marginally at the highest adult densities. The probability that the interaction was smaller than zero was 0.735. The slope of surface rugosity tilted from weakly positive at low adult abundance to weakly negative at high adult abundance. Still, the credible band overlapped zero across nearly the entire range, reflecting uncertainty in both the main effect and the interaction.

**Figure 3.**
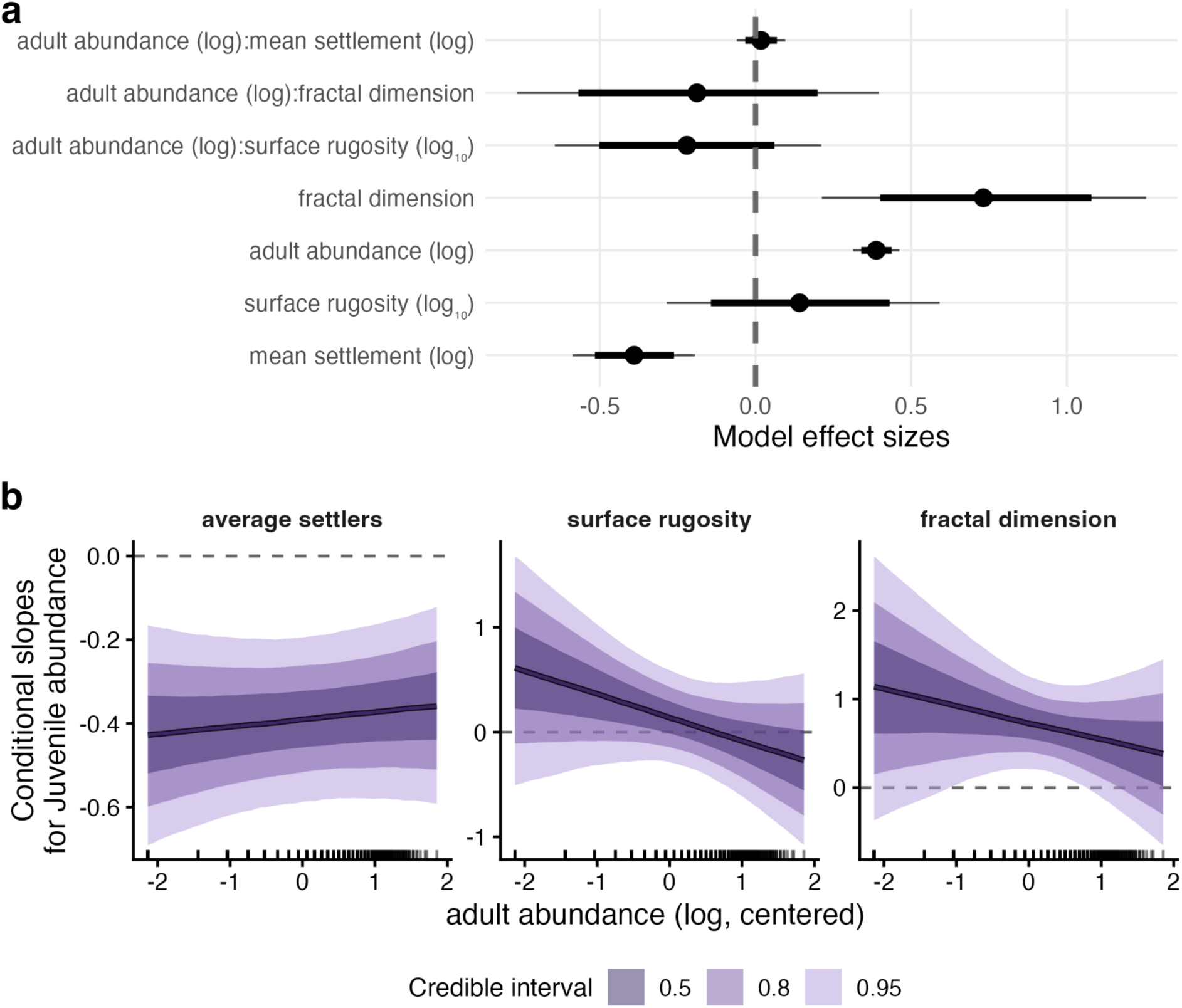
Adult abundance, reef fractal dimension, and settlement independently affect juvenile coral abundance, with some trends emerging in their interactions. (a) Posterior distributions of standardised regression coefficients from a Bayesian model of log-transformed juvenile abundance. Points indicate posterior medians; thick and thin lines show 80% and 95% credible intervals (CIs), respectively. Adult abundance (posterior mean = 0.39, 95% CI = [0.31, 0.46]) and fractal dimension (posterior slope = 0.73, 95% CI = [0.21, 1.25]) had credibly positive effects on juvenile abundance, while average settler abundance had a credibly negative effect (posterior mean = -0.39, 95% CI = [-0.59, -0.19]). Surface rugosity and all three interactions between adult abundance and habitat or settler predictors had 95% CIs overlapping zero, providing no credible evidence that adult abundance modifies these effects. (b) Conditional slopes of each predictor on juvenile abundance across the observed range of adult abundance. Solid lines indicate posterior medians; shaded ribbons show CIs. The marks on the x axis show where data were observed. The slope of average settler abundance remains negative, and that of fractal dimension remains positive, across essentially the entire range of adult abundance. In contrast, the slope of rugosity is poorly constrained, and its CIs encompass zero throughout. For comparison with other scales, refer to Figure SM8 and SM10

Juvenile abundance was strongly associated with local adult abundance at the 1 x 1 m scale (Figs. 3a and 4). The posterior mean effect of adult abundance was consistently positive (posterior mean = 0.39, 95% CI = [0.31, 0.46], Figs. 3a and 4). On the response scale, this corresponds to an approximately 48% increase in expected juvenile abundance for a one-unit increase in centred log adult abundance(Fig. 4). For example, increasing adult abundance from 10 to 20 adults was associated with an estimated 28.7% increase in expected juvenile abundance, while increasing from 10 to 11 adults was associated with an estimated 3.5% increase.

**Figure 4.**
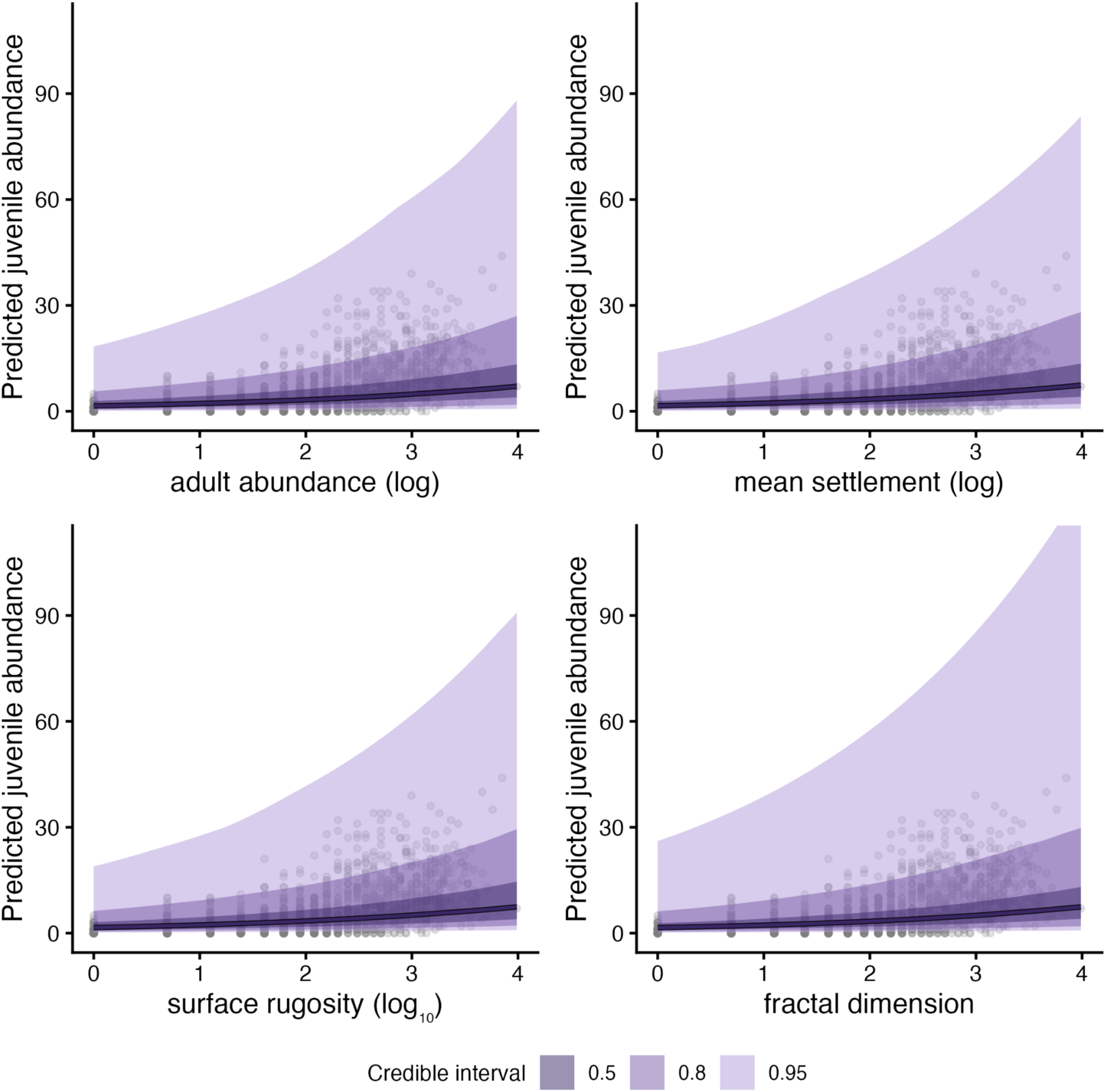
Adult abundance and fractal dimension positively affect juvenile abundance, while average settlement has a negative effect and surface rugosity has no detectable effect. Posterior expected marginal prediction of juvenile abundance based on each predictor used in the model. Solid lines indicate the median predicted value and shaded ribbons show 50%, 80% and 95% credible intervals (CIs). Dots show observed data. For comparison with other scale grids, refer to Figures SM11 and SM12.

Juvenile abundance also varied with settlement and habitat structure, independently (Figs. 3 and 4). Mean settlement was negatively associated with juvenile abundance (posterior mean = -0.39, 95% CI = [-0.59, -0.19]). In contrast, fractal dimension was positively associated with juvenile abundance, with a posterior mean of 0.73 (95% CI = [0.21, 1.25]). The effect of surface rugosity was more uncertain (posterior mean = 0.14, 95% CI = [-0.29, 0.59]).

Substantial variation in our response variable was attributable to site (standard deviation = 0.85, 95% CI = [0.56, 1.32]) and year (standard deviation = 1.80, 95% CI = [0.49, 5.25]), although the wider interval for year reflects the small number of levels (n = 2) of that factor (Fig. SM6). The negative binomial shape parameter was well-identified (φ = 2.83, 95% CI = [2.48, 3.22]), indicating moderate overdispersion relative to a Poisson distribution.

This pattern was largely robust to changes in spatial resolution (Fig. SM6). At the finer 0.5 x 0.5 m scale (Figs. SM8 and SM9), there was some evidence that the adult–juvenile association weakened with increasing fractal dimension, as indicated by a negative adult: fractal dimension interaction (posterior mean = -0.37, 95% CI = [-0.71, -0.03]). The adult-by-settlement and adult-by-rugosity interactions remained uncertain. At this finer scale, juvenile abundance also increased with adult abundance, with a posterior mean of 0.33 (95% CI = [0.28, 0.39]). Juvenile abundance also declined with mean settlement (posterior mean = -0.36, 95% CI = [-0.51, - 0.22]), and increased with fractal dimension (posterior mean = 0.37, 95% CI = [0.09, 0.64]). The effect of rugosity remained uncertain (posterior mean = 0.12, 95% CI = [-0.13, 0.38]).

At the coarser 2 x 2 m scale (Figs. SM10 and SM11), the direction of the main effects was also consistent with the 1 x 1 m analysis, although credible intervals were wider. Juvenile abundance increased with adult abundance (posterior mean = 0.46, 95% CI = [0.30, 0.61]) and with fractal dimension (posterior mean = 1.30, 95% CI = [0.19, 2.37]). At this scale, the estimated effects of mean settlement and surface rugosity were more uncertain, with credible intervals overlapping zero. Interactions between adult abundance and settlement, surface rugosity or fractal dimension were also uncertain. Overall, the multi-scale analyses indicate a robust positive association between local adult and juvenile abundance, together with a generally positive association with fractal dimension and limited evidence that habitat or settlement context strongly modified the adult–juvenile relationship.

## Discussion

We found that juvenile coral abundance, measured as the number of colonies, was associated with both local adult coral abundance and reef structural complexity, with effects acting largely additively within 1 × 1 m reef patches. Juvenile abundance was higher in reef patches with greater adult abundance and higher fractal dimension, whereas surface rugosity had an uncertain effect. Contrary to our initial expectation, tile-based settlement was negatively associated with juvenile abundance, suggesting that settlement and survival to the juvenile stage are decoupled. Overall, these patterns are consistent with coral assemblages influencing recruitment both directly, through adult-associated processes, and indirectly, through habitat structure. However, we found limited evidence that habitat complexity or settlement context strongly modified the adult–juvenile relationship. Thus, rather than detecting a strong synergistic or antagonistic interaction between adults and engineered reef habitat, our results point primarily to additive positive contributions of adult abundance and reef fractal dimension.

The positive association between adult and juvenile abundance suggests that post-settlement success is not independent of the local adult assemblage. Several non-exclusive mechanisms could generate this pattern. One possibility is facilitation^57^. Adult colonies might, for example, influence local consumer dynamics by diverting predation or grazing pressure away from small colonies ^58^. These mechanisms would be consistent with positive density dependence during early post-settlement stages. However, the same pattern could also arise through environmental filtering: sites or microhabitats that favour adult persistence might also favour juvenile survival, producing a positive adult–juvenile association without requiring direct facilitation by adults. Distinguishing facilitation from shared habitat filtering will require approaches that separate adult and juvenile survival trajectories over time and track them across gradients of reef complexities.

Structural complexity also contributed to juvenile abundance, but not all dimensions of complexity behaved similarly. Fractal dimension was consistently positively associated with juvenile abundance across spatial scales, whereas surface rugosity showed weaker and uncertain effects. This distinction suggests that the aspects of reef structure most relevant to early coral survival do not simply reflect total surface area. Fractal dimension better captures fine-scale branching, crevices, and other geometric irregularities across multiple spatial scales^47^, potentially providing a more sensitive measure of the structural complexity relevant to coral recruitment and the availability of settlement microhabitats. Surface rugosity, by contrast, might describe broader elevation differences without necessarily discriminating across the specific structural features used by settlers and juveniles. This result supports the idea that reef complexity is multidimensional, and that different metrics capture different ecological functions^42,47,59^.

Conditional effects suggested that fractal dimension has stronger positive effects at lower adult abundance, but this pattern was uncertain at the main 1 m scale and should be interpreted cautiously. At this scale, none of the interactions between adult abundance and the habitat or settlement predictors was credibly different from zero. However, the finer 0.5 x 0.5 m analysis provided some evidence of antagonistic effects of fractal dimension on juveniles with increasing adult abundance, suggesting that a buffering mechanism could be in place. One possible interpretation is that fine-scale structural complexity can partially buffer low adult abundance by providing refugia, settlement surfaces, or favourable microsites. Under this scenario, complexity would contribute more where adult abundance is low, reducing juvenile abundance’s dependence on nearby adults. More broadly, the multi-scale analyses indicate that adult abundance is the most robust predictor of juvenile abundance. At the same time, fractal dimension provides an additional structural signal whose strength might depend on the spatial grain at which recruitment processes are measured.

Our results should be interpreted in light of the fact that adult coral abundance was estimated using colony counts rather than colony cover, biomass, or size-based metrics. Counts provide a consistent measure of colony presence and numerical abundance, but they do not capture variation in colony size or three-dimensional space occupation. As a result, the interaction patterns we detect could differ if we measured adult abundance as coral cover or accounted for size structure, particularly if larger colonies contribute disproportionately to habitat structure, fecundity, or competitive effects. For example, patterns could differ if adult corals were weighted by planar cover or size structure, because a reef with many small colonies could function differently from one with fewer but larger colonies^60,61^. However, alternative metrics would also introduce their own limitations. Planar cover estimates derived from orthomosaics may underestimate true coral abundance by excluding colonies present on vertical surfaces, under overhangs, or within structurally complex parts of the reef ^62^. In our case, this is particularly relevant because up to a third of colonies were present in the reef but not visible in the maps. Thus, while using counts may affect the strength or interpretation of relationships involving reef structure and recruitment, cover- or size-based approaches would not necessarily provide an unbiased alternative. Future work combining colony counts with size and/or cover would help clarify how robust the adult–recruit relationship we found is to different representations of adult abundance.

Our findings point to the importance of ecological inheritance in coral reef recovery. After disturbances such as bleaching, dead coral skeletons can continue to shape the geometric structure of the reef even after live coral cover declines. This inherited structure can create microhabitats that support juvenile survival and, in turn, influence the trajectory of community recovery. In our study system, where sites experienced severe disturbance that led to mass mortality^63^ and were previously considered to have low recovery potential due to limited larval supply^64^, the positive association between reef fractal dimension and juvenile abundance suggests that the physical legacy of past coral assemblages continues to benefit subsequent generations^65^. Similar results have been found for different life-stages in corals^66^ and in other systems, like oyster reefs^15^. Our results are consistent with a niche-construction feedback in which populations generate habitat features that can later influence recruitment and persistence^66,67^.

The negative association between tile-based settlement and juvenile abundance was unexpected. Settlement tiles are widely used to estimate larval supply and reef recovery potential, but our results add to evidence that tile settlement and *in situ* juvenile abundance do not always align^10,68^. Several mechanisms could explain this mismatch. First, the taxa settling on tiles could differ from those later detected as juveniles on natural reef substrata. Second, early post-settlement mortality mechanisms, such as crowding effects from density dependence, could also occur at the assemblage level but went undetected in our field observations because they act before coral colonies were visible as juveniles ^27^. Third, the juveniles observed in benthic surveys might include individuals that settled in previous years and only became visible after a time lag, creating a “seed bank” effect of cryptic recruits^30^. If so, within-year settlement estimates from tiles will not correspond closely to visible juvenile abundance in the same survey period, as also found in another analysis from the same sites^68^. Finally, settlement tiles remove the natural structural and biological context of the reef, including adult neighbours, cryptic refugia, and fine-scale surface heterogeneity. As a result, tile-based estimates capture larval arrival or initial settlement rate, but operate on a time scale too short to reflect post-settlement survival processes that shape juvenile abundance on natural reef surfaces. Our results reinforce that early life-stage post-settlement processes can play a larger role in early coral survival than previously appreciated.

The scale-sensitivity analysis further suggests that density-dependent and structural effects operate across spatial grains, but with different levels of uncertainty. At the 0.5 m scale, adult abundance, tile-based settlement, and fractal dimension showed patterns broadly consistent with the 1 m analysis, and evidence suggested that complexity could modify the adult–juvenile relationship. At the 2 m scale, adult abundance and fractal dimension remained positively associated. Still, uncertainty increased for several predictors, likely reflecting fewer spatial units and coarser aggregation of fine-scale recruitment processes. These results indicate that the positive adult–juvenile association is robust. In contrast, habitat complexity is more sensitive to the spatial scale at which reef structure and recruitment are quantified. This is expected if microhabitat conditions at centimetre-to-metre scales strongly influence early coral survival.

From a management perspective, our findings can help design long-term strategic actions for reef protection. Structural complexity alone increases settlement rate ^26,66^, and we showed that structure can also promote successful coral juvenile recruitment. But our findings also show that juvenile recruitment increases with live adult colony abundance, suggesting that focusing on maintaining live adult colonies could contribute not only to replenishing coral populations but also to maintaining higher recruitment rates and fine reef structural complexity.

Niche construction mechanisms might depend on the life stage of the organisms experiencing them, with the benefits adults gain from modifying their environment differing from the benefits those modifications provide to incoming recruits. Here, we found that adult populations and structural complexity independently predicted juvenile abundance, with little evidence of interaction, suggesting that both the coral assemblage and the associated structural habitat shape post-settlement recruitment. Protecting and restoring reef structure, alongside efforts like adult outplanting, might be a valuable, independent lever for supporting recruitment. More broadly, these results suggest that ecosystem engineers’ legacy persists and supports recruitment independently of the engineers themselves, offering a mechanism by which reefs can retain recovery potential even as adult populations decline.

## Acknowledgements

We thank the Australian Museum’s Lizard Island Research Station staff for their support, Marion Chapeu for her help in the field, and Fabio Guzzo for his help with the conceptual figure. This research was supported by: a John Templeton Foundation grant (grant #60501 ’Putting the Extended Evolutionary Synthesis to the Test’, awardees: Maria Dornelas and Joshua S Madin); a MAST small grant (SG396, awardee: Viviana Brambilla); a Leverhulme fellowship (awardee: Maria Dornelas); three Ian Potter Doctoral Fellowships at Lizard Island Research Station (awardees: Viviana Brambilla, Damaris Torres-Pulliza, Devynn Wulstein); an Isobel Bennett Marine Biology Fellowship (awardee: Nina Schiettekatte); a University of Hawai’i start-up fund (awardee: Joshua S Madin); a Charles Warman Foundation Lizard Island Research Foundation Critical Grant ‘Understanding coral reef recovery from extreme disturbances using 3D maps’ (awardees: Maria Dornelas and Joshua S Madin); a National Science Foundation–Natural Environment Research Council Biological Oceanography Grant (grant 1948946, awardees: Joshua S Madin and Maria Dornelas); an ERC-CoG (CoralINT, GA 101044975, awardee: Maria Dornelas). Funding was provided by the European Union. Views and opinions expressed are however those of the authors only and do not necessarily reflect those of the European Union or the European Research Council. Neither the European Union nor the granting authority can be held responsible for them.

## Supplementary material

**Figure SM1.**
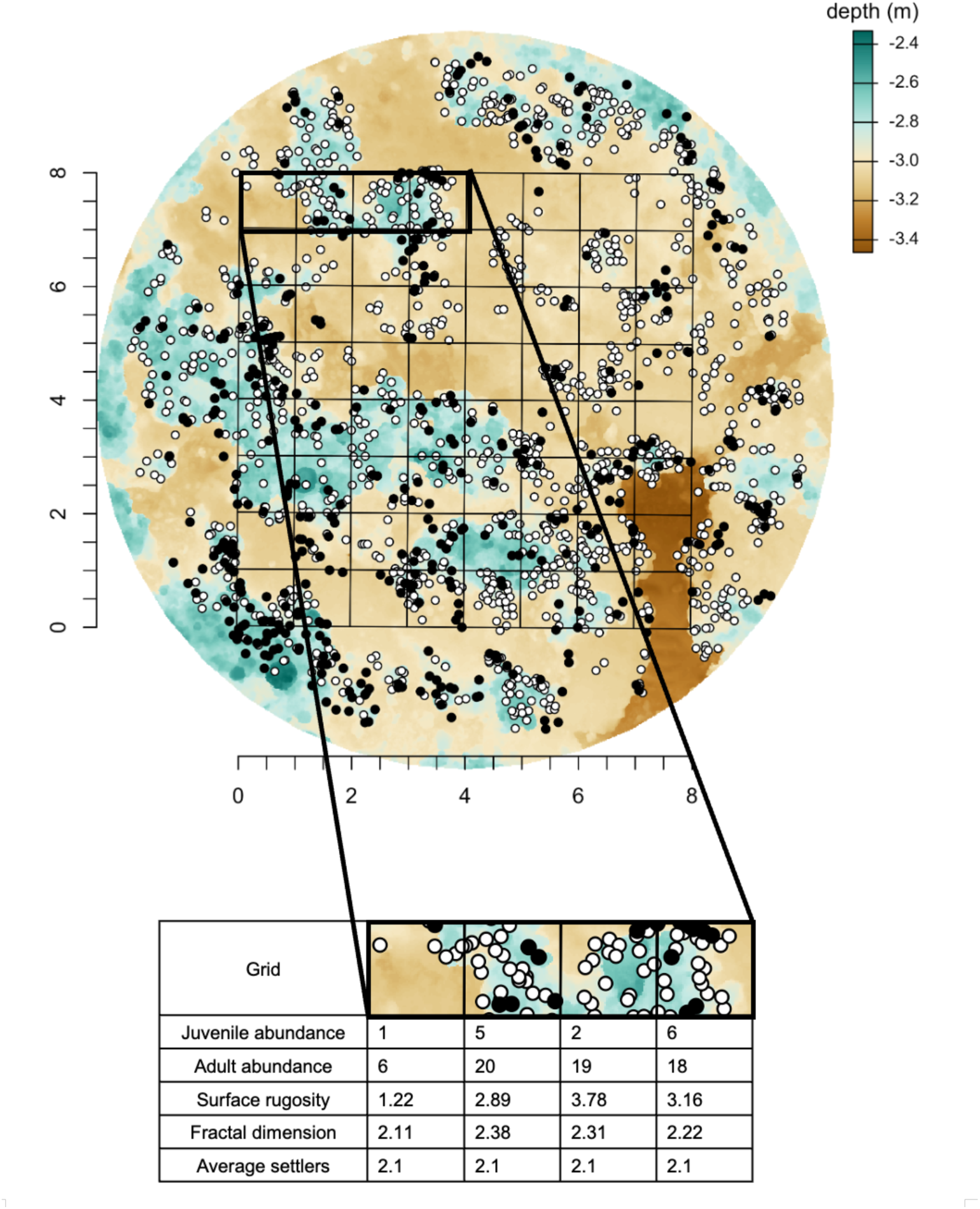
Digital elevation model and spatially explicit coral occurrence dataset used to estimate complexity and count adults and juveniles during each sampling event in each quadrat. Digital elevation model of the reef site at North Reef 3 (N3) in 2019 with the black 1 × 1 m grid overlaid and variable per grid examples. Black dots represent the locations of juvenile coral colonies, and white dots represent the locations of adults. Both juvenile and adult dots include both cryptic and hidden colonies that were only visible in the field upon careful inspection. Notice that the average settler abundance is constant, as there was only one settlement estimate per reef site per sampling event.

**SM2.**
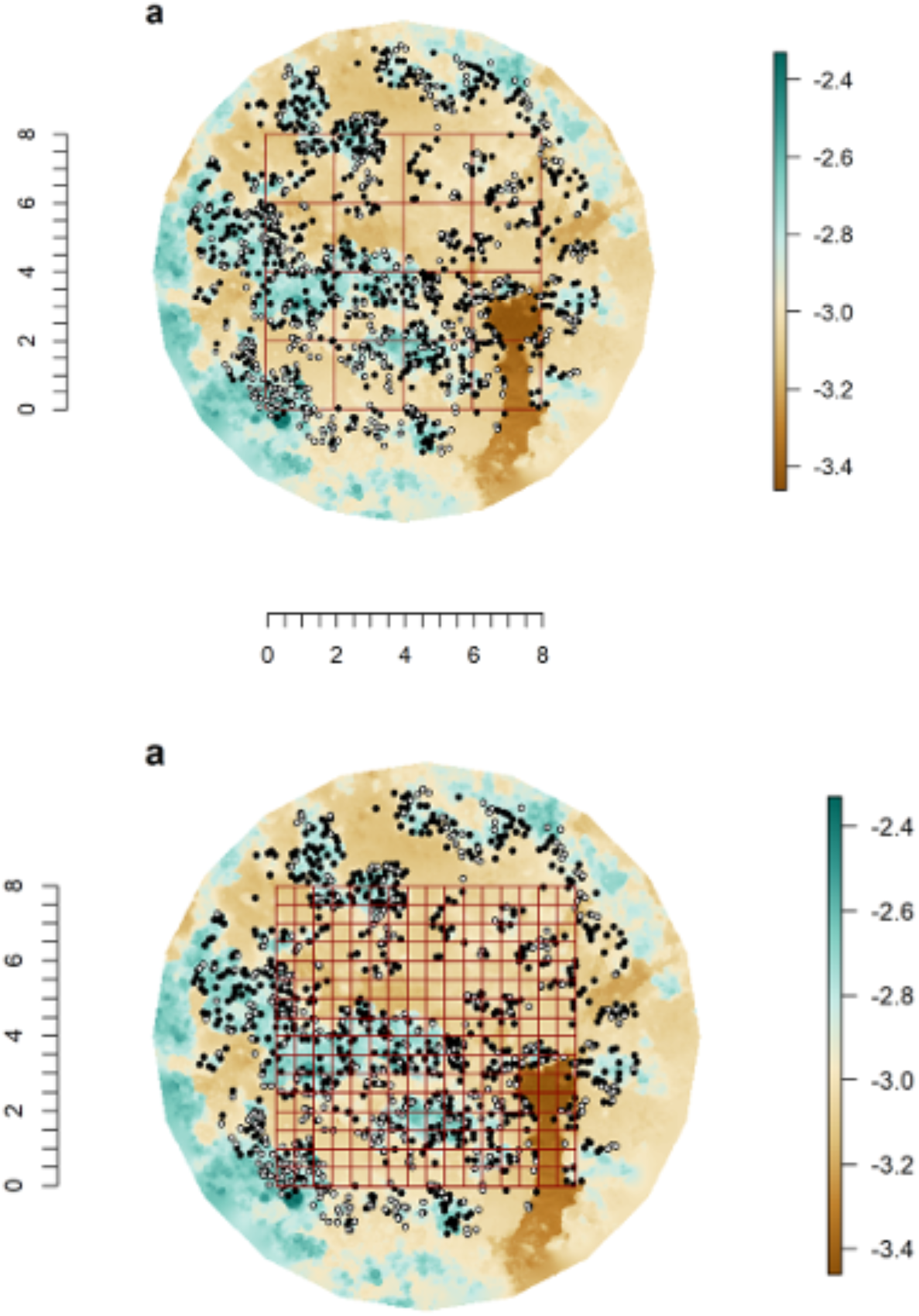
Digital elevation model example and juvenile and adult colonies for the 0.5 x 0.5 m grid and 2 x 2m grid. Digital elevation model of the reef site at North Reef 3 (N3). Black dots represent the locations of adult colonies and white dots represent the locations of juveniles. In dark red, the grids that divide across (a) the 0.5 x 0.5 m scale, resulting in 256 quadrats of area 0.25 m^2^ per site (n = 5376 in total), and (b) 2 x 2m scale, resulting in 16 quadrats of area 4 m^2^ per site (n = 336 in total). Colour represents depth.

**SM3.**
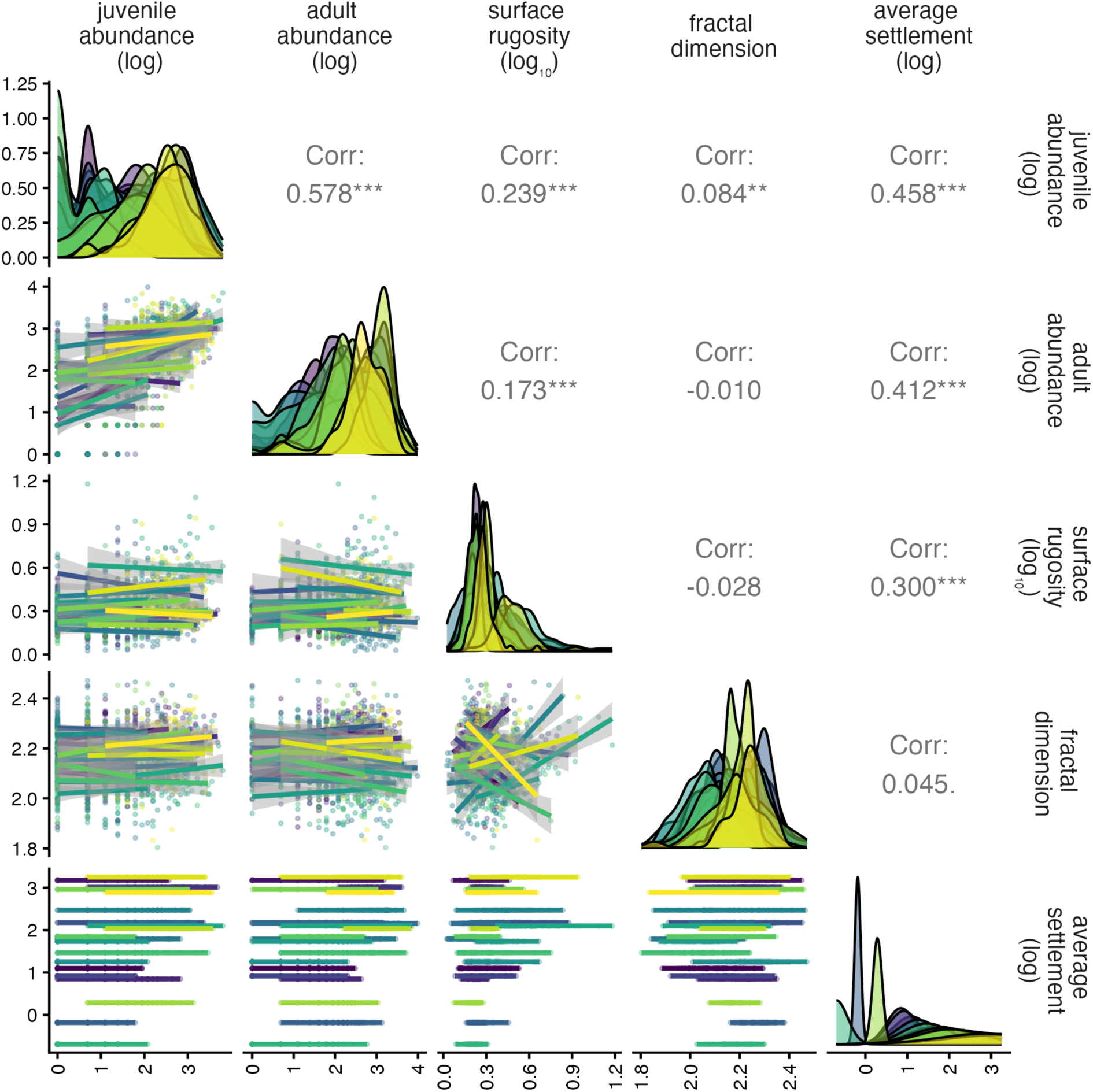
Pairplot of variables measuder in each quadrat at the 1 x 1m grid. Colour represents different sites.

**Figure SM4.**
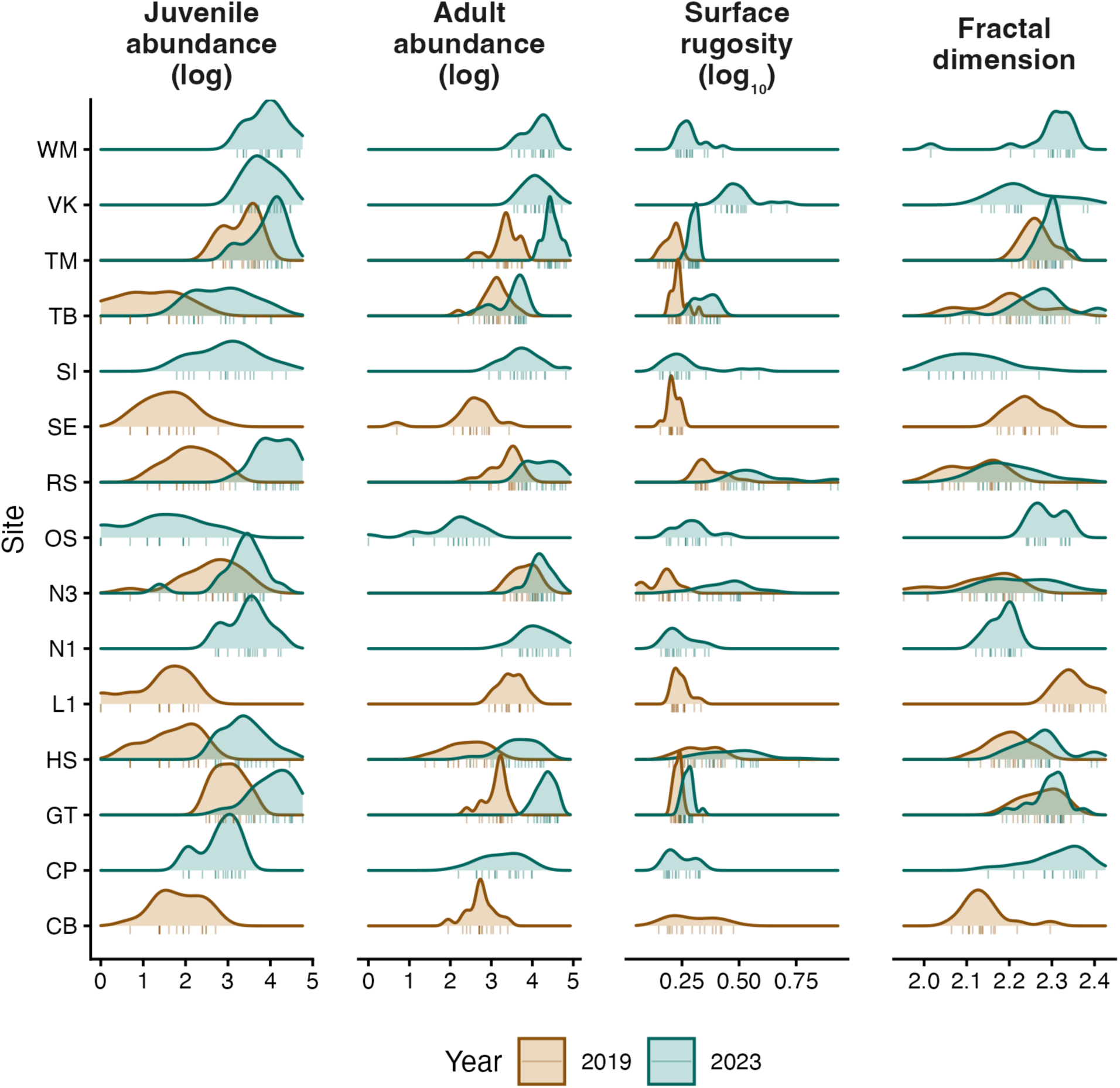
Density distribution of the variables for bigger scale sensitivity analysis (2 x 2m scale) shows overall variation across sites and years sampled (n = 366).

**Figure SM5.**
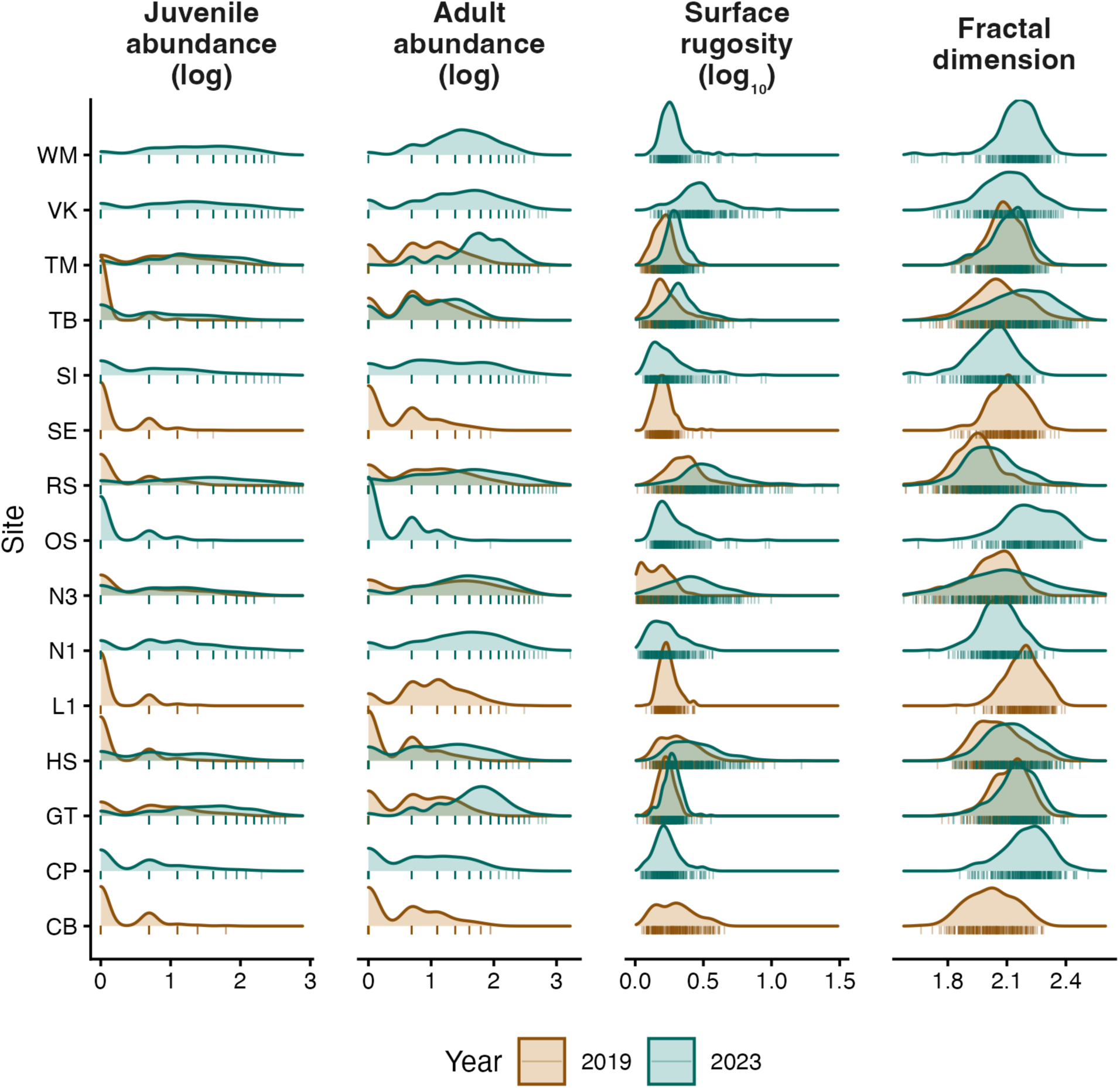
Density distribution of the variables for smaller scale sensitivity (0.5 x 0.5 m scale) shows overall variation across sites and years sampled (n = 5376).

**Figure SM6.**
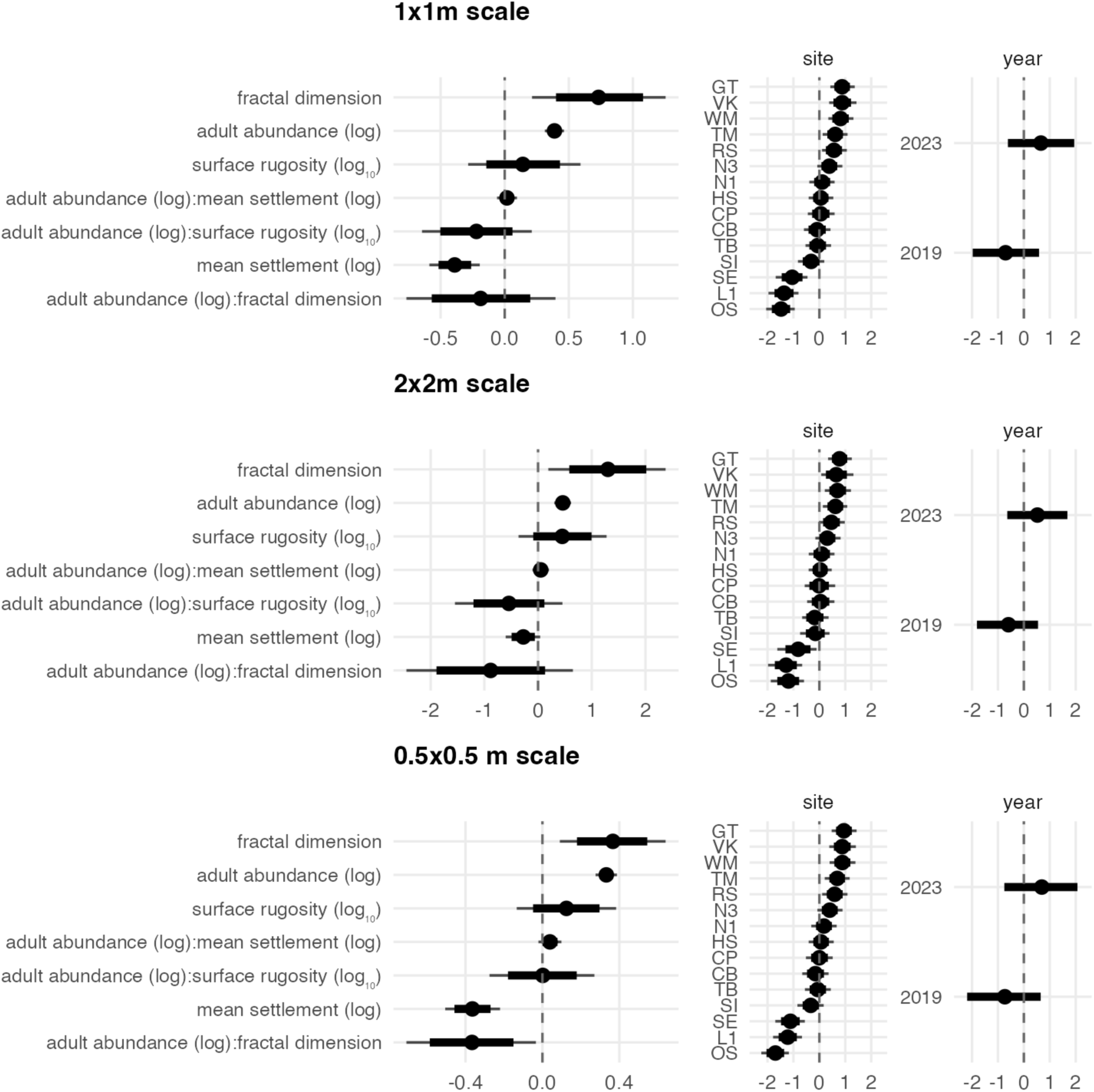
Fixed and random effect sizes across scales. (a) Posterior distributions of standardised regression coefficients from a Bayesian model of log-transformed juvenile abundance. Points indicate posterior medians; thick and thin lines show 80% and 95% credible intervals (CIs), respectively.

**Fig SM7.**
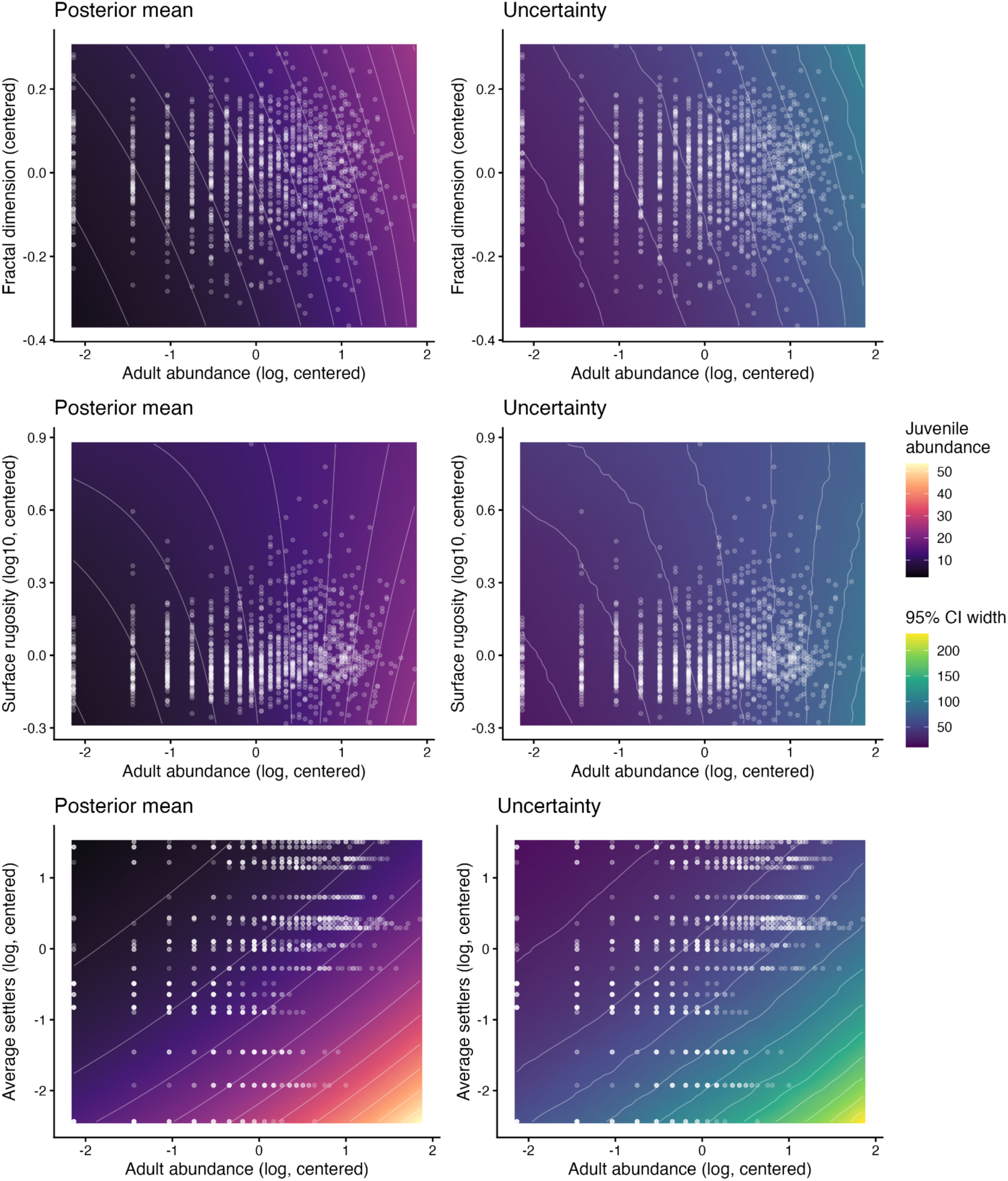
Predicted coral recruitment across variable pairs show little interaction effects at the 1 x 1 m scale. On the left, posterior mean prediction heatmaps. On the right, uncertainty is given by the width of the CIs. Notice that variables have been centred. Dots represent data pairs used to fit the model.

**Fig SM8.**
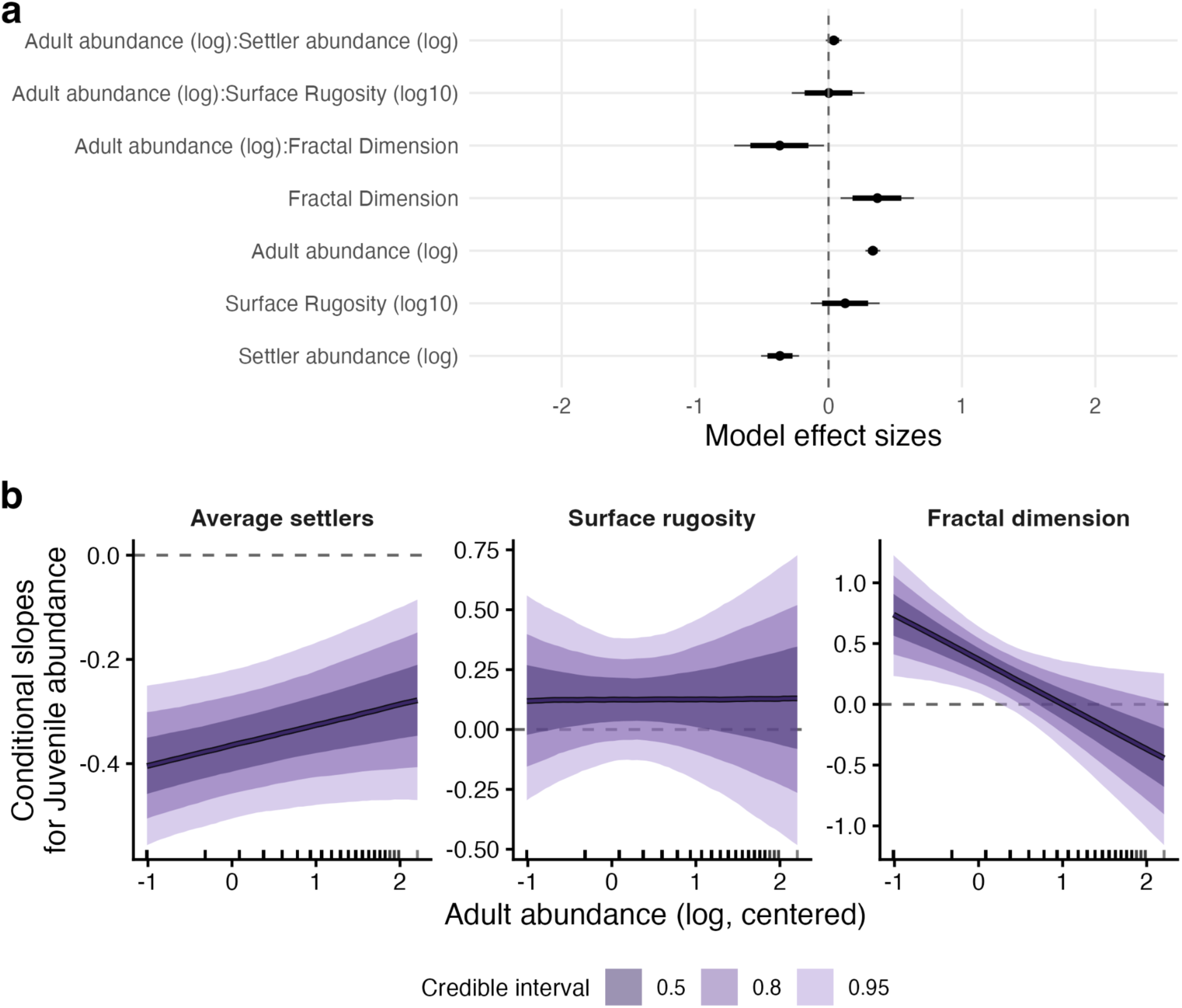
Adult abundance and fractal dimension have a negative interaction effect, and settlement has an independent effect on juvenile coral abundance at the 0.5 × 0.5 m scale. (a) Posterior distributions of standardised regression coefficients from a Bayesian model of log-transformed juvenile abundance. Points indicate posterior medians; thick and thin lines show 80% and 95% credible intervals (CIs), respectively. Adult abundance and fractal dimension had a negative interaction and positive effects on juvenile abundance, while average settler abundance had a credibly negative effect. Surface rugosity and the interactions between adult abundance and rugosity or settler predictors had 95% CIs overlapping zero, providing no credible evidence for these effects. (b) Conditional slopes of each predictor on juvenile abundance across the observed range of adult abundance. Solid lines indicate posterior medians; shaded ribbons show CIs. The marks on the x axis show where data were observed. Slopes are poorly constrained, but the slope of fractal dimension decreases along the range of adult abundance. Dashed horizontal lines mark a slope of zero.

**Fig SM9.**
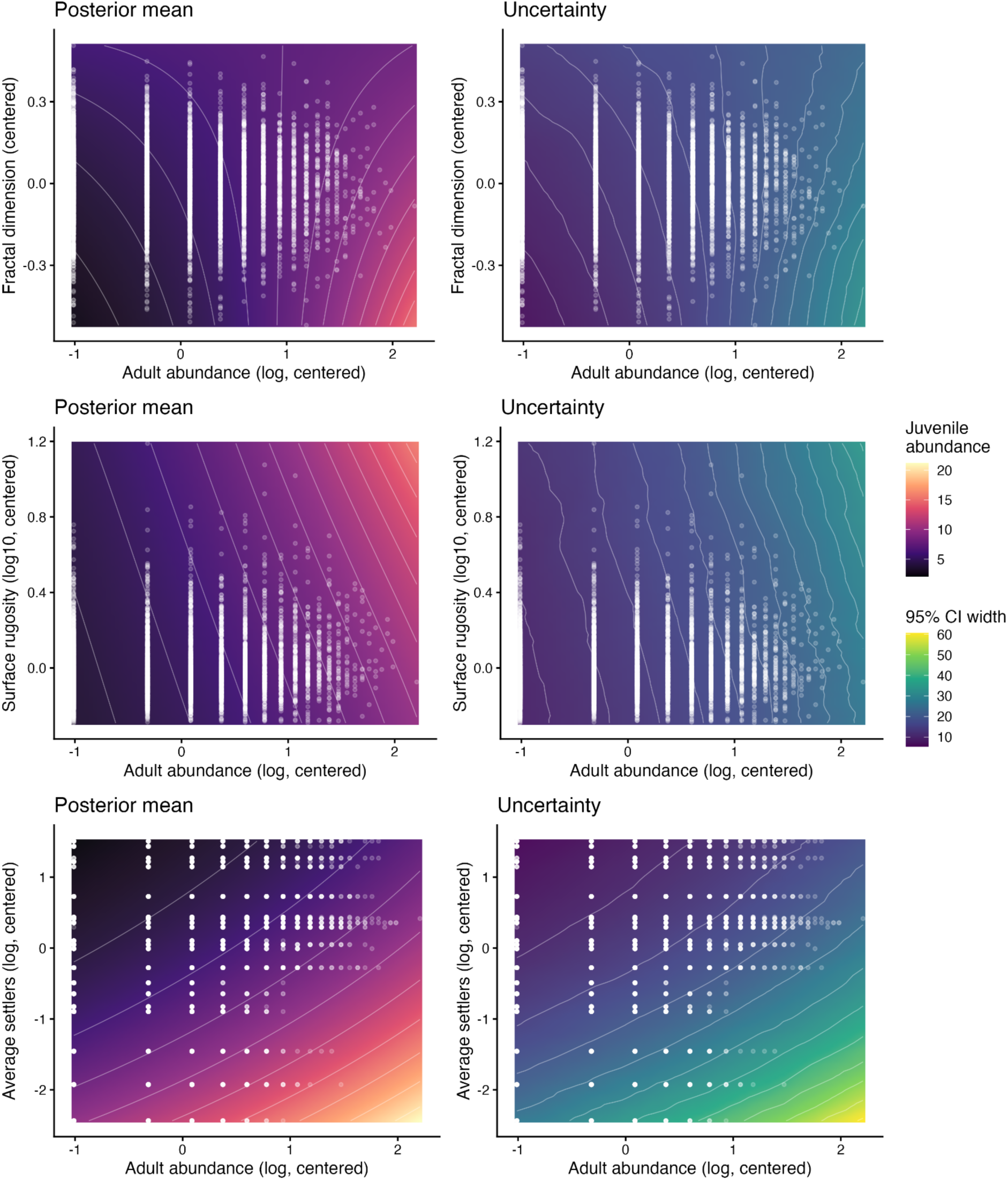
Predicted coral recruitment across variable pairs show little interaction effects at the 0.5 x 0.5 m scale. On the left, posterior means prediction heatmaps. On the right, uncertainty is given by the width of the CIs. Notice that variables have been centred. Dots represent data pairs used to fit the model.

**Fig SM10.**
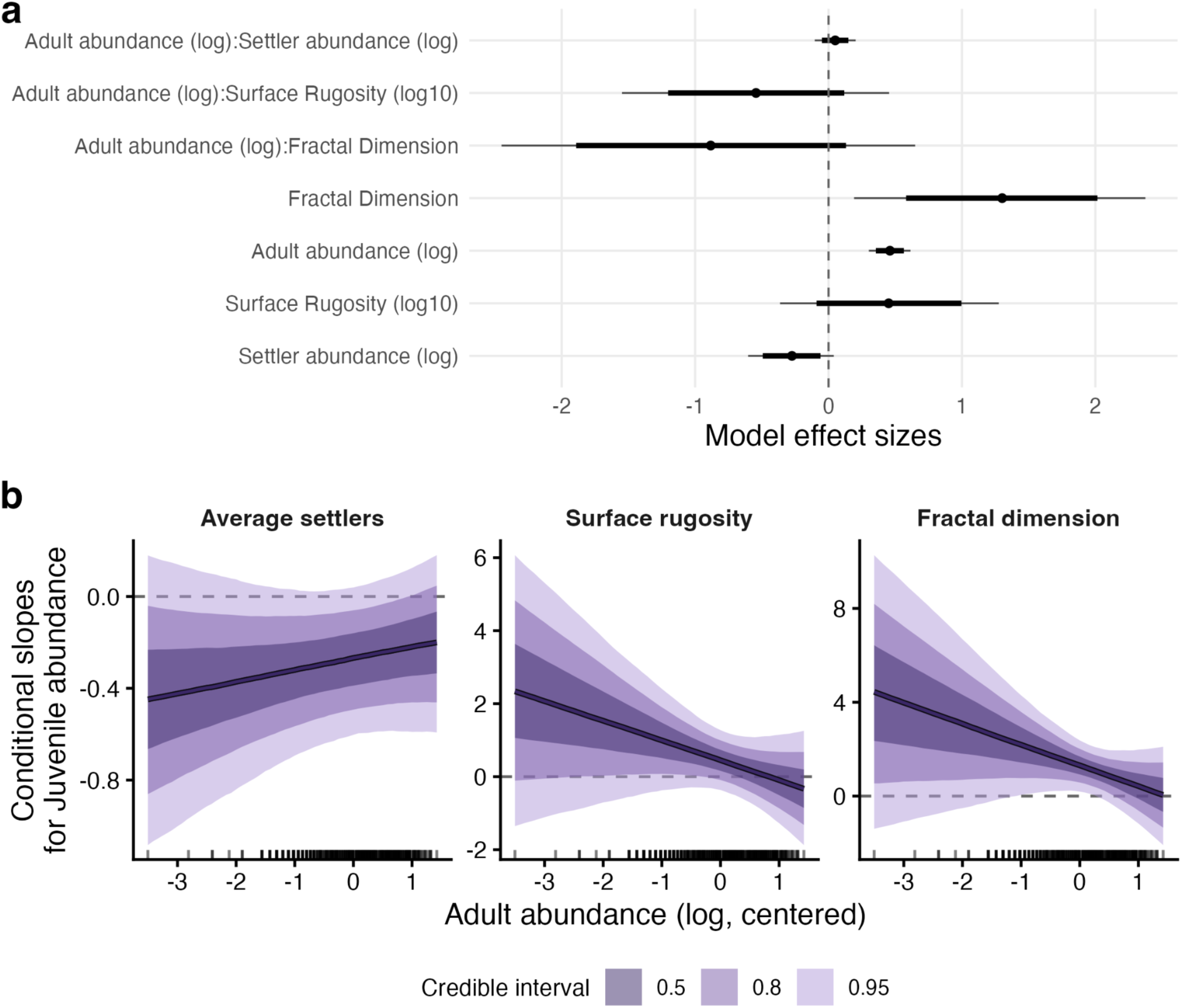
Adult abundance and habitat structural complexity have independent effects on juvenile coral abundance at the 2 × 2 m scale. (a) Posterior distributions of standardised regression coefficients from a Bayesian model of log-transformed juvenile abundance. Points indicate posterior medians; thick and thin lines show 80% and 95% credible intervals (CIs), respectively. Adult abundance and fractal dimension had credibly positive effects on juvenile abundance, while average settler abundance had a credibly negative effect. Surface rugosity and all three interactions between adult abundance and habitat or settler predictors had 95% CIs overlapping zero, providing no credible evidence that adult abundance modifies these effects. (b) Conditional slopes of each predictor on juvenile abundance across the observed range of adult abundance. Solid lines indicate posterior medians; shaded ribbons show CIs. Marks on the x-axis show where data were observed. The slope of average settler abundance remains negative, and that of fractal dimension remains positive, across essentially the entire range of adult abundance. In contrast, the slope of rugosity is poorly constrained, and its CI encompasses zero throughout. Dashed horizontal lines mark a slope of zero.

**Fig SM11.**
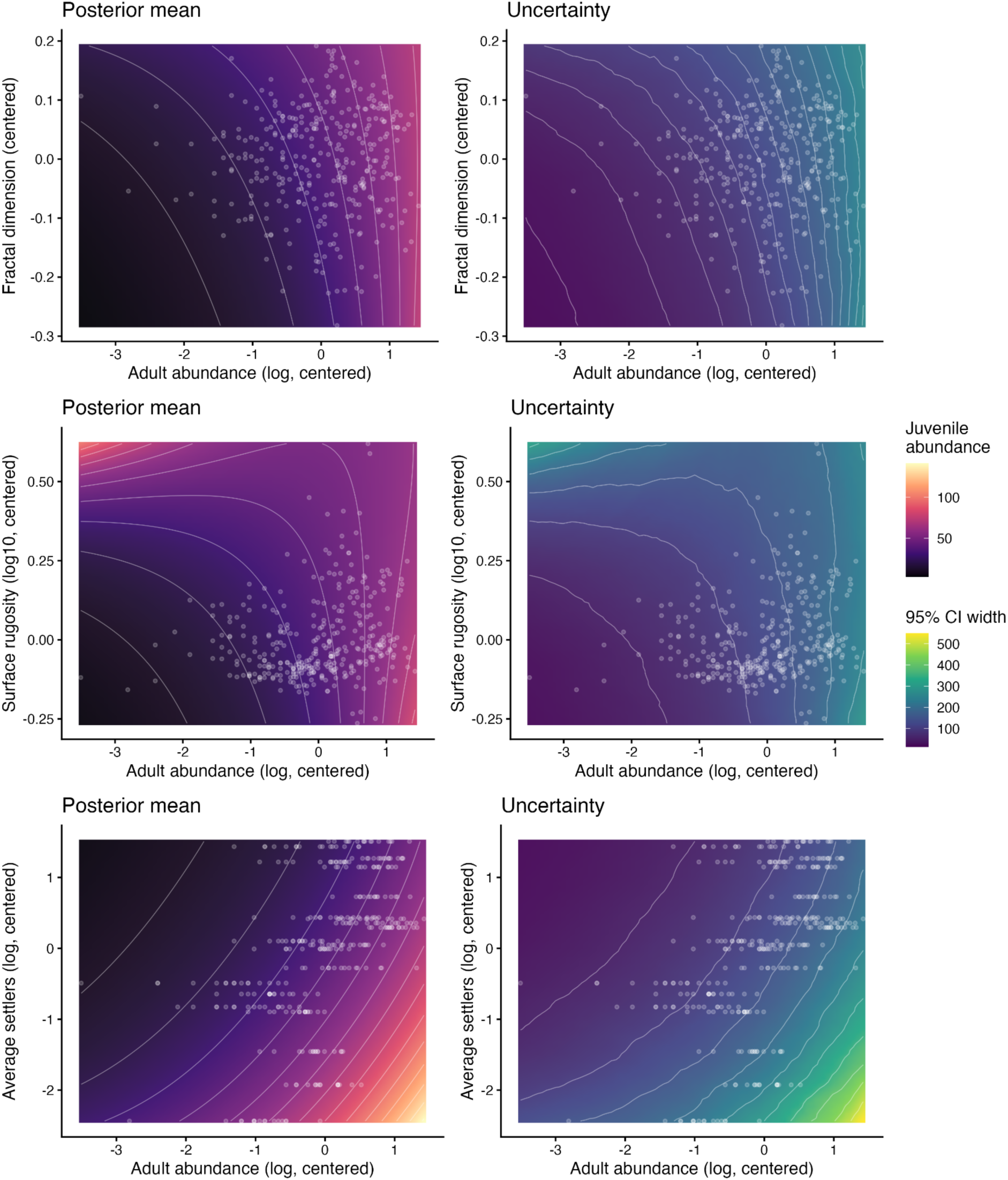
Predicted coral recruitment across variable pairs show little interaction effects at the 2 x 2 m scale. On the left, posterior mean prediction heatmaps. On the right, uncertainty is given by the width of the CIs. Notice that variables have been centred. Dots represent data pairs used to fit the model.

